# Learning and Increased Effort Affect Lipid Gene Ezxpression in the Mushroom Bodies of Bombus terrestris; A Possible Model for Flower Handling?

**DOI:** 10.1101/2025.06.25.661662

**Authors:** Kalliu Carvalho Couto, Rafael F. da Silva, Linnea A. M. Erlingsson, Eirik Søvik, Johannes Gjerstad

**Affiliations:** Faculty of Health Sciences, Oslo Metropolitan University, Oslo, Norway; Mental Health Services, Akershus University Hospital, Lørenskog, Norway; Volda University College, Volda, Norway; Kristiania University College, Faculty of Health Sciences, Oslo, Norway

**Keywords:** bumblebee, learning, flower-handling, mushroom bodies, mRNA sequencing, gene expression

## Abstract

**Objective:** In this study, we examine the effects of instrumental learning, and increased effort on the gene expression profile in the mushroom bodies of bumblebees’ (Bombus terrestris).

**Methods:** In a low-effort group (n=9), Bombus terrestris learned to press a lever to access sugar water solution (instrumental task) and responded with a constant effort for a period of four hours, whereas in a high-effort group (n=10), subjects were trained with the same task, but effort requirement was doubled in the last two hours of the experiment. The gene expression profiles of these two groups were compared with a control group (n=10) where subjects had access to the sucrose solution ad libitum, and no learning or effort manipulation took place. After running the experiment, total RNA from each bumblebee’s mushroom bodies was shipped to Novogene for high-throughput RNA sequencing (mRNAseq). Gene Ontology (GO) analyses were performed to evaluate the effects of the two training protocols on biochemical pathways. Regression analyses were used to examine the relationship between the top five mRNAs and bumblebee performance on high- and low-effort tasks.

**Results:** In total, 150 and 112 genes were deregulated by the low- and high-effort procedures, respectively. Complete hierarchic clustering regarding low- and high-effort groups versus the control was observed. GO analyses of the low- and high effort groups versus control revealed that the deregulated genes were associated with pathways related to lipid metabolism. Analyses of the top five deregulated genes suggested a link between mRNA of LOC100646091 to performance following low effort protocol.

**Conclusion:** The data from the present study showed that exposure to the low- and high-effort reinforcement procedures may be associated with the deregulation of many genes related to lipid metabolism in the bumblebee’s mushroom bodies. Additionally, data suggest a link between expression of mRNA of LOC100646091 and performance. The procedure used here—learning of an operant task and an increase in the effort required (number of lever presses)—is proposed as a model for bumblebee flower-handling.

## Background

Bumblebee foraging depends on several complex and interdependent behavioural repertoires. For example, when searching for nectar and pollen, bumblebees need to navigate between flower patches, learn and remember flowers’ sensorial characteristics as well as allocate foraging duration and the frequency of visitation to different flowers (Dukas & Visscher, 1994). Recent data show that octopamine signalling in the bee brain affects the chemosensory responsiveness (reward sensitivity) and therefore also decision-making in honeybee foragers (Arenas et al., 2021). How bee foragers cope with complexity in needs and resources is, however, a major question in behavioural ecology (Francis et al., 2016).

Several lines of evidence show that foraging alters gene expression profile of the mushroom bodies important for learning and memory (Kamikouchi et al., 1998; Lutz et al., 2012; McNeill et al., 2016). Some findings also indicate that the gene expression profile of the mushroom bodies could reflect social behaviour including communication between individuals (Sen Sarma et al., 2009). Moreover, gene expression changes after colour learning may involve several transcriptional large-scale waves suggesting a molecular mechanism of long-term memory formation (Li et al., 2018). Regarding olfactory learning, many genes may be differently expressed between learner bees and failed-learner bees (Raza et al., 2022).

At the cellular level, the foraging behaviour driven by reward may be linked to several key molecules. These include the neuromodulators such as dopamine and octopamine (Agarwal et al., 2011; Huang et al., 2022; Peso et al., 2016). Although the details of how these molecules affect (or are affected by) the cellular reward signalling remains unclear, evidence exists that many so-called immediate early genes (genes that control other genes) are deregulated by foraging flight (Iino et al., 2020). Also, earlier observations suggest that pheromones alter expression patterns of foraging-related genes in the bee’s brain (Ma et al., 2019).

Most previous experimental studies of the gene expression employed a proboscis extension reflex (PER) protocol to evaluated involuntary reflexive responses. PER studies use a classical conditioning procedure in which bees are immobilized, and a neutral stimulus (visual or olfactory) is paired with an unconditioned stimulus (US; sugar water) for a number of trials. If learning occurs, the conditioned stimulus (CS) will elicit the PER. Delayed exposure to the CS is used to evaluate memory acquisition (Huang et al., 2023) or visual stimuli i.e., colour training (Li et al., 2018). Although necessary for successful foraging, the cognitive functions that are assessed in the PER assay are insufficient. During foraging, bumblebees must be able to handle flowers to access pollen and nectar. Flowers that are visited by bumblebees are known to exhibit a wide range of morphological diversity, varying in complexity and the amount of effort that is required to access their nectar and pollen (Laverty et al., 1988). Therefore, the ability to handle different types of flowers is a critical component of successful foraging in bumblebees. Handling flowers requires a high level of cognitive function, and impairments in this activity have been suggested to decrease foraging efficiency (Phelps et al., 2020). Bumblebees must learn and remember how to handle flowers to access floral rewards, which involves problem-solving activities such as pushing, pulling, and twisting floral structures. They also need to adjust their handling strategies to different floral morphologies, which requires learning flexibility.

Against this backdrop, the present study adopt an operant conditioning approach to investigate the link between genetic profiles and instrumental learning and effort level, proposing a bumblebee animal model for flower handling. This flower-handling model involves: 1) the learning of a complex task, in the form of lever press responses to obtain sucrose solution, and 2) the manipulation of effort by increasing the number of lever responses required to access the reward. This procedure measures bumblebees’ performance in a task requiring complex instrumental learning (i.e., collecting syrup by activating an operandum) and effort manipulation (i.e., varying the number of lever presses required). For this purpose, subjects were trained to fly across a flying arena and land on an apparatus where lever responses were learned through shaping successive approximations, and the requirement to operate the dispenser (number of lever presses) was manipulated. Effort-related schedules that manipulate response requirements have been extensively used to study the effects of toxicants and brain alterations on learning and motivation in other species. For example, studies have demonstrated that rats depleted of dopamine had their performance affected in such tasks (Salamone et al., 2012). However, no study has been performed on instrumental learning, and increased effort on the gene expression profile of bumblebees.

## Method

### Subjects

All bumblebees (*B. terrestris*, naïve, unknown age) used in the present study were obtained from the same farm in Norway (Bombus AS). Upon arrival, pollen was continuously available inside the colonies, which were kept in a room with at a temperature of ~25 ^0^C, at a relative humidity of ~45%. All experiments were conducted in the light period of an artificial 12 h light/12 h dark cycle.

### Apparatus

An adapted version of an operant chamber first described by Pessotti (1969) and later automated and modified according to the instructions of Escobar and Santillán (2017) and Penha et al. (2024), was used. The operant chamber consisted of transparent an acrylic box (30 cm x 30 cm x 30 cm) covered with a grey 3D-printed surface, placed inside the wooden box. One of the walls featured a lever that controlled the delivery of sugar-water from the opposite wall (figure 1). All experimental events were controlled and recorded using a program in Visual Basic Express 2010 and Arduino IDE.

**Figure 1.**
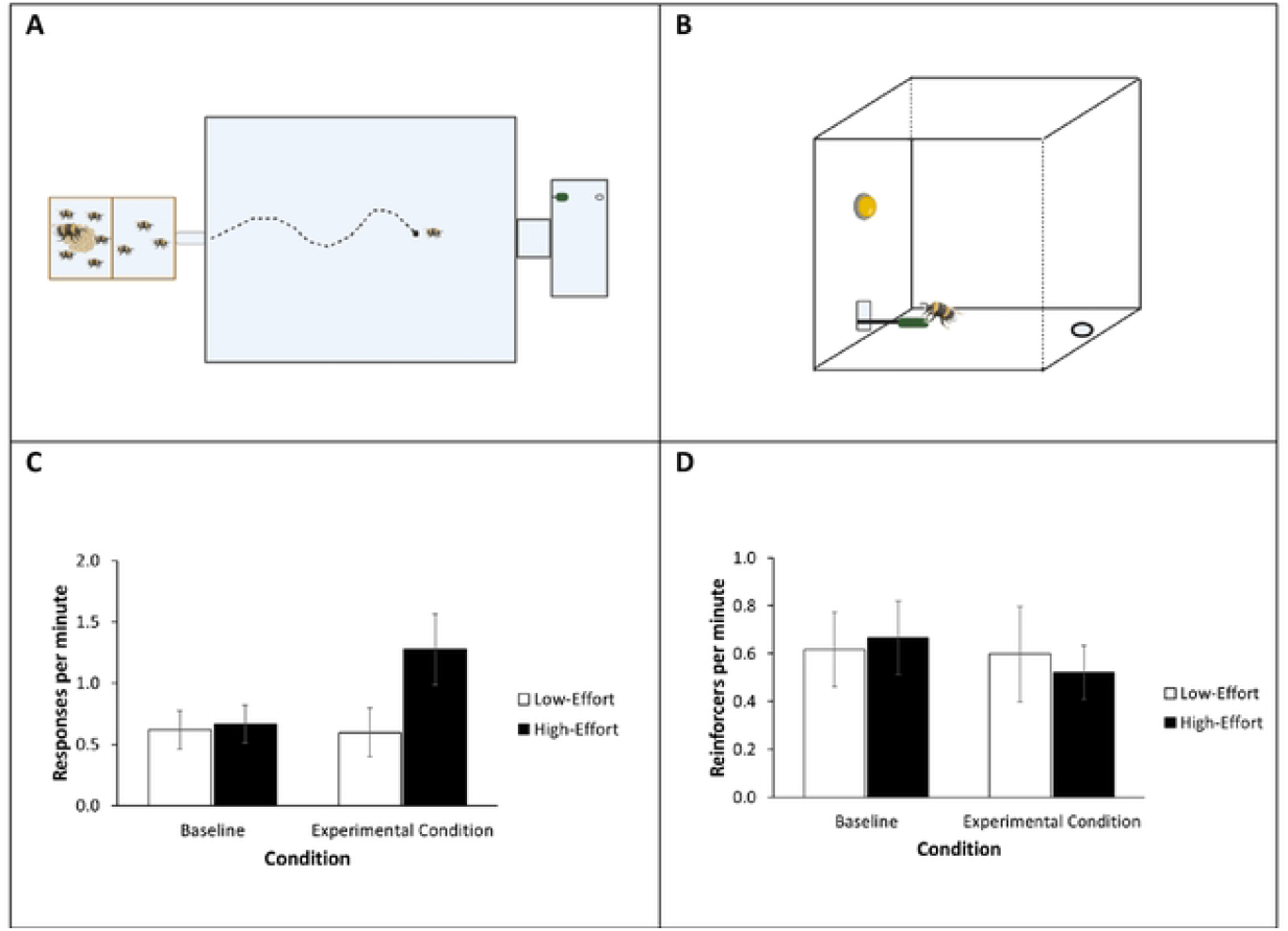
Panels A and B provide a visual representation of the experimental set-up, while panels Cand D present the data on responses and reinforcements per minute. Panel A depicts the flying arena where the colony is connected. Bumblebees fly across this arena to reach the operant chamber. Panel B shows the operant chamber where the bumblebees perform the bar-pressing task to obtain reinforcement. Panel C shows the responses per minute (y axis) of bumblebees subjected to low-effort (white bars) and high-effort (black bars) groups during baseline and experimental conditions. Panel D presents the reinforcement rate (y axis) of bumblebees subjected to low-effort (white bars) and high effort (black bars) groups during baseline and experimental conditions

### Pre-training – Flying arena and Lever Pressing Shaping

Before the experimental sessions, the foraging bumblebees in each colony underwent training to enter the flying arena and reach operant chamber (Figure 1A and 1B). Subjects that reached the operant chamber apparatus were picked up and marked with different numbers glut on their backs.

Experimental sessions began with a selection process in which a small cup was placed in the center of the operant chamber, and bumblebees could feed freely. In each session the first subject to complete five visits (i.e., land, feed, and return to the colony) was selected for the study. Any subject that reached the apparatus after the selection was removed and placed in a waiting box separated from the flying arena.

Selection was followed by lever-pressing shaping procedure, consisting of five reinforced steps: (a) feed from the feeder, (b) turn toward the lever, (c) get close to the lever, (d) step on the lever and (e) press it, separated by several visits to the apparatus.

For each step, the first responses in the first three visits were cued by a small drop of sugar-water solution, and once the subject learned where the target behaviour, no more cues were provided and target behaviour was reinforced in the apparatus feeder for three more visits.

### Experimental conditions

In total, 28 bumblebees were selected for 28 days and included in the study. Three groups were used to examine the association between 1) learning and 2) level of effort to changes in the global gene expression profile. The groups consisted of; Low Effort (L) and High Effort (H) procedures, where the bees were trained to reach and press a lever to obtain sugar-water solution, in addition to Control (C) procedure were bumblebees flew to the apparatus and sugar water was available ad libitum.

Subject in the *Low Effort (L) group* were exposed learning procedure, where lever pressing responses were shaped through differential reinforcement of successive responses, until lever pressing was learned, followed by a two hours baselined in Fixed Ratio (FR) schedule of reinforcement that required one response to activate the feeder (FR1), and experimental condition with two more hours of FR1 exposure. Each lever-press response activated the feeder once (1:1), making sugar water solution was available for 15 seconds, signalized by an LED light. Thus, in this condition, subjects were learned an instrumental task, but effort (response requirement) was maintained constant across baseline and experimental condition at one lever press.

Subjects in the *High Effort (H) group* were also exposed learning procedure, where lever pressing responses were shaped through differential reinforcement of successive responses until learned, followed by a two hours FR1 baseline condition. However, effort was manipulated during the two hours in the experimental condition, by increase the effort requirement to an average of two responses to every reinforcers delivered (2:1). For that, subjects were exposed to an variable ratio (VR) schedule of reinforcement where the feeder was activated after an average of two responses (VR2) to (signalized by an LED light), ranging between 1, 2, and 3 responses. Thus, subject in condition learned an instrumental task, and level of effort was manipulated during the condition two.

In *Control (C) group*, bumblebees were free feeding i.e., non-learning or effort manipulation. Subjects of this group went through only the pre-training in the flying arena, and four hours of feeding in the apparatus with no lever presses requirement.

### Tissue harvesting, Dissection and RNA Isolation

After the going through the experimental protocol, the bumblebees were captured, and their brains were dissected, for more details see (Carreck et al., 2013). The trachea, the hypopharyngeal glands, the salivary glands, the retinal pigment as well as the optical lobes were removed. The remaining parts of the brains i.e., the mushroom bodies, were then snap frozen on dry ice and stored in –80 °C. According to the manufacturer’s protocols, the frozen tissues were later homogenized by a pestle in a mortar, and total RNA isolated using the PureLink™ RNA Mini Kit (Thermo Fisher Scientific Inc., USA).

Intact total RNA was isolated from the 28 subjects. Individual RNA from all bumblebees in the Low effort group (n=9), High effort group (n=9) and Control group (n=10), were shipped to Novogene for RNA sequencing (mRNAseq), for shipping details see (Olsen et al., 2022).

### RNA sequencing (RNAseq) and analyses

Novogene performed quality control and cleaning of the raw RNAseq reads, and mapped the clean reads to the rat reference genome (Rnor_6.0, Ensembl release 80) (Cunningham et al., 2021) by using HISAT2 (Kim et al., 2019). DESeq2 was used by Novogene to obtain differentially expressed genes (DEGs), where significance was accepted at p < 0.05 (Love et al., 2014). Further genetic analyses were conducted online on Novogene’s e-Portal (Customer Service System). The Benjamini-Hochberg procedure was applied to control for false discovery rate FDR at a 5% (Padj = 0.05). Gene Ontology (GO) analyses of the low and high effort groups versus the control group were used to examine the effect of the training protocols. Regression analyses were used to examine the association between top three mRNA-fold expression and response rates.

## Results

Regarding the responses per minute of bumblebees subjected to low-effort and high-effort conditions across baseline and experimental phases, the low-effort group is represented with white bars and the high-effort group with black bars (Figure 1C and 1D). During the baseline condition, the response rates were similar between the two groups, with the low-effort group averaging 0.61 responses per minute (SEM = 0.06) and the high-effort group averaging 0.66 responses per minute (SEM = 0.06). Over the two-hour experimental condition period, the response rates for the high-effort group doubled, reaching 1.26 responses per minute (SEM = 0.11), while the response rates for the low-effort group remained similar to baseline at approximately 0.56 responses per minute (SEM = 0.06). This outcome was expected, as the response requirement for the high-effort group in the experimental condition was on average of twice the number of responses for reinforcer (2:1 ratio; with response requirement varying between 1, 2, and 3) when compared to baseline, while the low-effort group remained responding to an FR1 schedule. Although response rates for high-effort group doubled during the experimental condition compared to the low-effort group, the reinforcement rate was slightly higher for the first 0.66 (SEM = 0.06) when compared to the later 0.61 (SD = 0.06).

The RNAseq yielded an average of 48.4 mill clean reads, of which 97.0% were mapped and 95.0% were uniquely mapped to the *B. terrestris* reference genome iyBomTerr1.2 (NCBI). The reads uniquely mapped genomes were used in the further gene expression analyses. Based on clean reads uniquely mapped to the reference genome, a total of 12 910 expressed genes were identified.

Regarding the three groups—low effort, high effort, and control —no clear clusters of the aggregated principal components 1 and 2 (PC1 and PC2) of the global gene expression (i.e., RNAseq) were observed (Figure 2). Still, the RNAseq analyses of each of the low- and high-effort groups versus the control group revealed that the RNA profiles of the mushroom bodies following the low-effort and high-effort procedures were different from the controls (Figure 3A and 3B). In total, 150 and 120 genes were deregulated within the low- and high-effort groups, respectively (40/29 up and 110/83 down). Thus, more genes were down-regulated than up-regulated during the two protocols where learning (low- and high-effort groups) and effort (high-effort) were manipulated.

**Figure 2.**
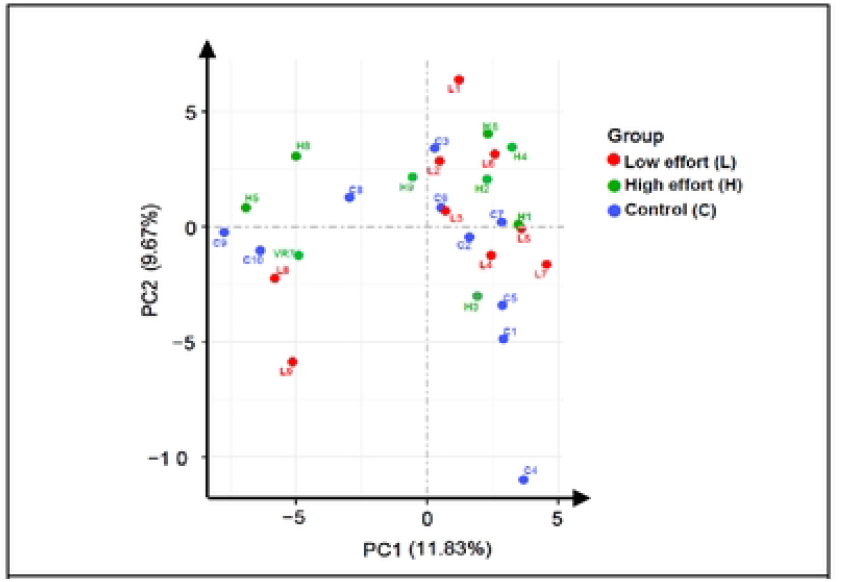
Principal component analysis (PCA) plot; the data demonstrate the distribution of gene expression profiles of the three groups low effort reinforcement (red), high effort reinforcement (green) and control (blue). The percentage of variance is indicated on the axes, with PC1 accounting for 11.83% and PC2 accounting for 9.67% of the total variance. No clear clusters of the aggregated principal components 1 and 2 of the global gene expression were observed.

**Figure 3.**
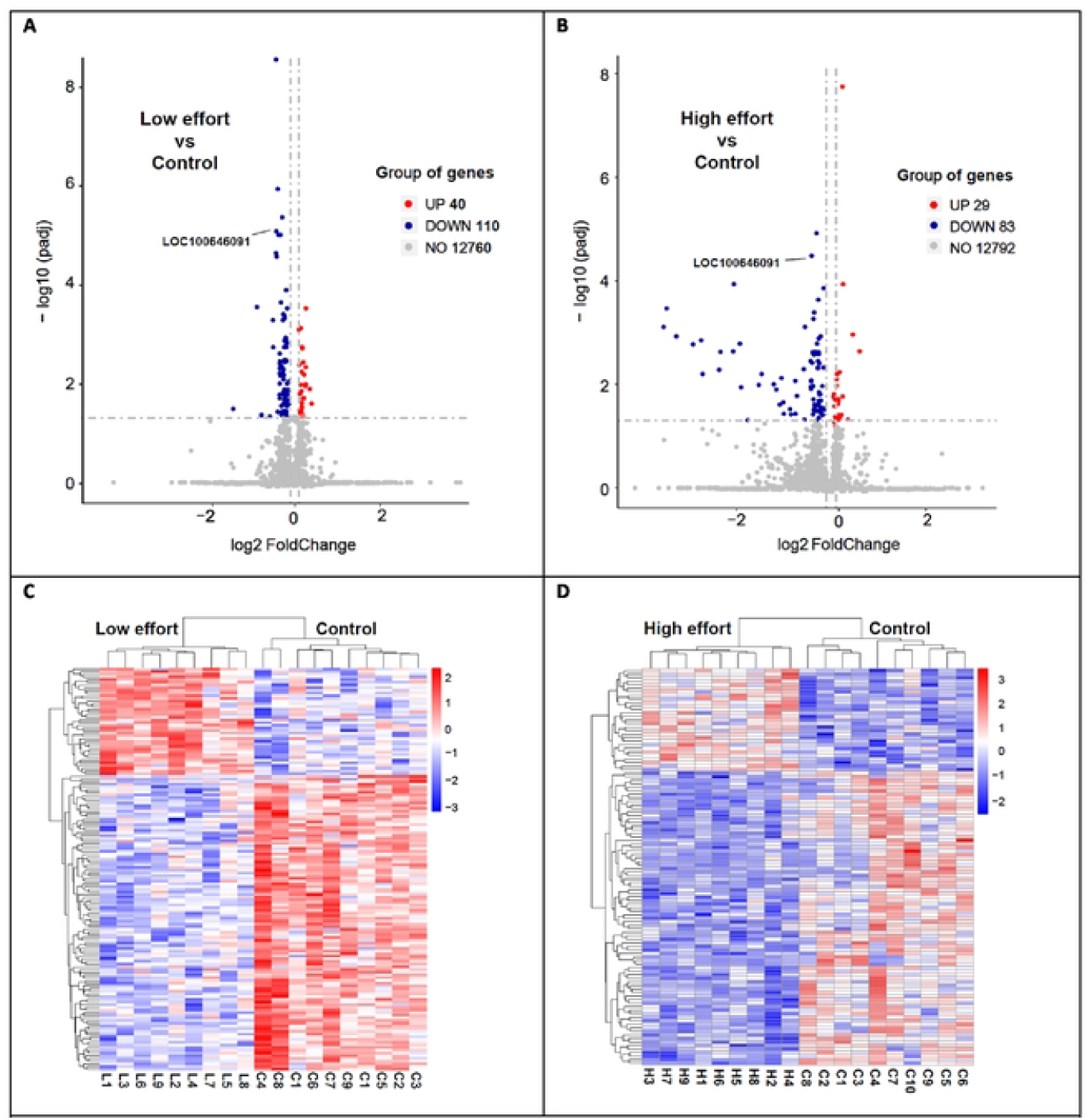
Effetct of training procedures on the brain bee gene expression. A and BJ Significance level and fold change of RNAseq gene expression in the low- or high effort groups, both n=9, relative to control i.e., the repetitive group, n=10. The data displayed 40/29 up-regulated mRNA (red dots) and 110/83 down-regulated mRNA (blue dots). Padj = 0.05 (Benjamini-Hochberg), fold change > 0.1. The gene LOC100646091 that appeared on top five in both analyses (Padj) are highlighted with its gene name. C and D) Heatmap of the 40/29 plus 110/83 mRNA significantly up- or down regulated genes by the learning procedures versus the control. The deepest red indicates the strongest up-regulation, whereas the deepest blue indicates the strongest down-regulation(normalized to the average of gene expression of all 150/112 mRNAs). Note the clustering; gene expression data of the bees in the low- or high effort groups are seen on the left of the gene expression data of the controls.

The RNA profiles of the low-effort and high-effort groups compared to the control group, showed complete hierarchical clustering (Figure 3C and 3D). Without exception, the RNA profiles of subjects within each group were more similar to one another than to the RNA profiles of any subject in the control group. Moreover, Gene Ontology (GO) analyses of each of the low- and high-effort groups compared to the control group revealed that the deregulated genes were associated with lipid metabolic or biosynthetic processes (figure 4A and 4B).

**Figure 4.**
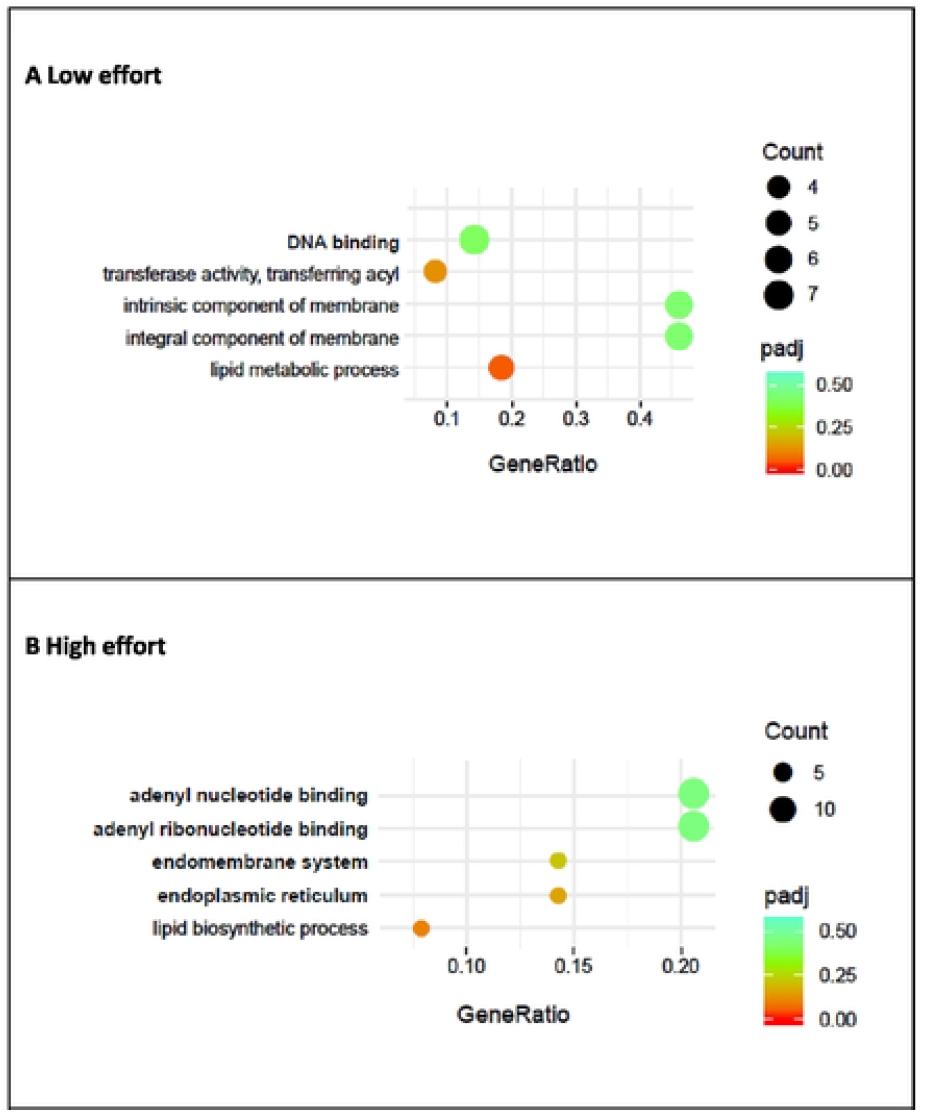
Clusters of genes affected by the different training procedures on the brain bee. A and B) Signalling pathways i.e., GO-terms affected by the different training procedures on the brain bee. Dot plot of top 5 enriched GO-terms produced after low effort and high effort reinforcement based on all significant deregulated genes. X-axis shows gene ratio, i.e., number of genes clustered to a given GO term divided by total number of genes in input list. Size of dots represents number of genes in the clusters. Colour green to red represents adjusted p-value of the GO-terms.

Listing the top five (Padj) differentially expressed genes (DEGs) in the low- and high-effort groups compared to the control group showed that the gene LOC100646091 appeared on both lists (table 1). However, additional analyses of the relationship between the top five mRNA expression levels versus responses, linked LOC100646091 to lever presses performance (measure as rates of responding) in the low-effort group (figure 5A), but not in the high-effort group (figure 5B).

**Figure 5.**
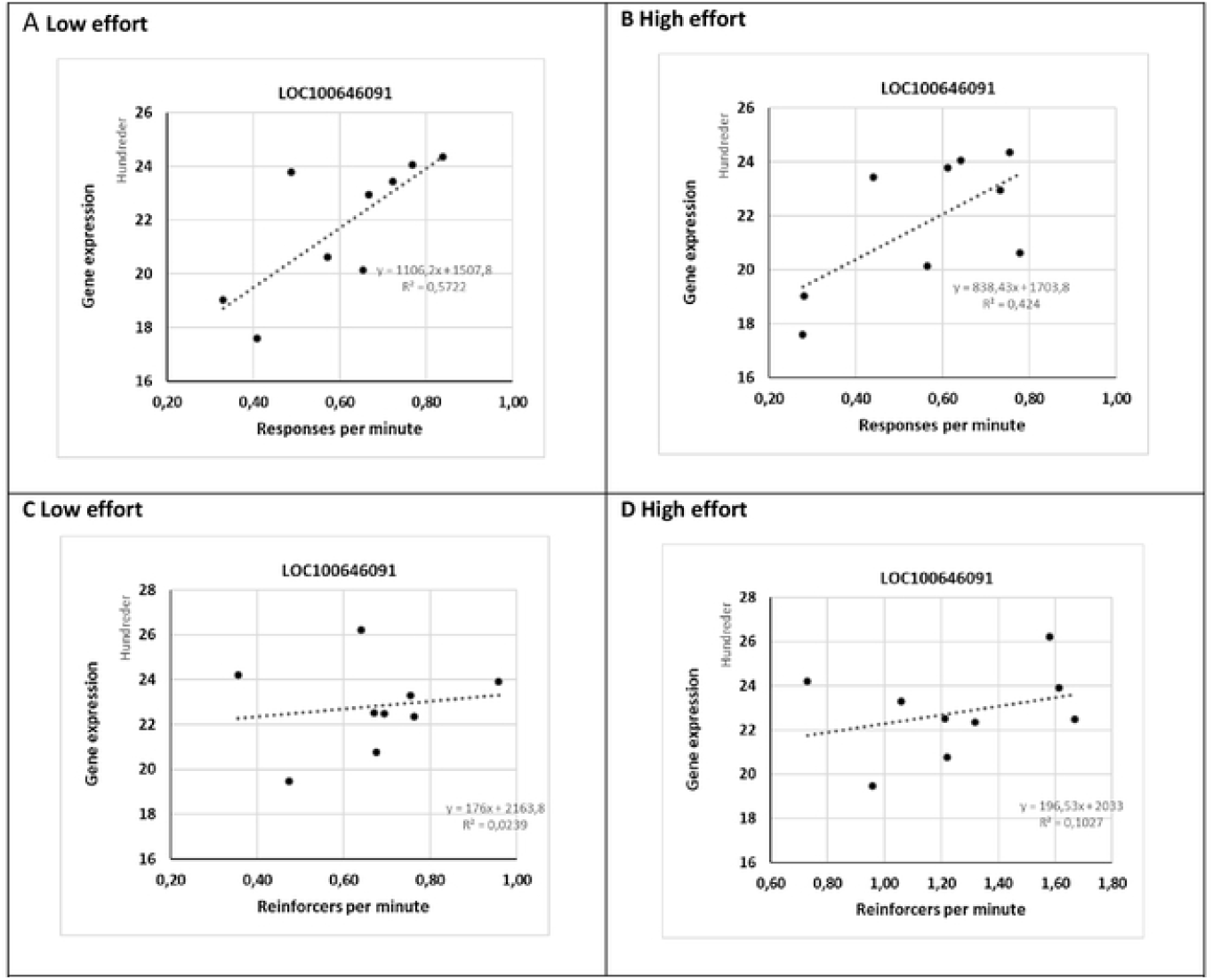
Regression analysis of the gene expression mRNA LOC100646091 versus performance (response rate) data of the 9 individual bees that underwent the A and B low effort (p=0.018/0.057) and Cand D the high effort (p=n.s./n.s.) training procedure, respectively.

**Table 1.** Top five differentiany expressed genes (DEGS) ranked by adjusted p values (padj) in low effort reinforcement (L) and high effort reinforcement (H) groups relative to the control (C) group. The gene expression of LOC100646091 (seen on both top five lists) was in the low effort reinf Drcement group associated with the perfomance, i.e., response rate (regression, p=0.018).

**Low effort reinforcement (L vs C)**
| Gene name | log2 Fold change | Padj CR vs Rep | Gene chr | Start | End | Biotype | Reg p |
| --- | --- | --- | --- | --- | --- | --- | --- |
| LOC100644393 | -0,4565 | 2,83E-09 | NC063271,1 | 14527964 | 14535448 | protein coding | n.s |
| LOC100652309 | -0,4228 | 1,18E-06 | NC063270,1 | 11554303 | 11562041 | protein coding | n.s |
| LOC105666697 | -0,3033 | 4,36E-06 | NC063286,1 | 3874896 | 3880073 | protein coding | n.s |
| **LOC100646091** | **-0,4572** | 8,34E-06 | NC063269,1 | 801446 | 805840 | protein coding | **0,018** |
| LOC100645127 | -0,3027 | 1,01E-05 | NC063276,1 | 2181312 | 2184740 | protein coding | n.s |

**High effort reinforcement (H vs C)**
| Gene name | log2 Fold change | Padj VR vs Rep | Gene chr | Start | End | Biotype | Reg p |
| --- | --- | --- | --- | --- | --- | --- | --- |
| LOC100650615 | 0,2448 | 1,76E-08 | NC_063280,1 | 8943387 | 8946683 | protein coding | n.s |
| LOC100643999 | -0,3110 | 1,22E-05 | NC_063269,1 | 4224890 | 4230428 | protein coding | n.s |
| **LOC100646091** | **-0,4075** | 3,30E-05 | NC_063269,1 | 801446 | 805840 | protein coding | n.s |
| LOC100647220 | -2,0611 | 0,0001165 | NC_063275,1 | 22716839 | 22719937 | protein coding | n.s |
| LOC100648299 | 0,2594 | 0,0001165 | NC_063282,1 | 10336456 | 10342396 | protein coding | n.s |

A broader approach listing the top 15 (Padj) differentially expressed genes (DEGs) in the low- and high-effort groups versus the control group showed an overlap of only four genes; LOC100646091, LOC125385672, LOC100648299 and LOC100643999 (Supplementary table 1).

## Discussion

In present study we show that learning procedures relevant for foraging affect both performance and gene expression in the **mushroom bodies** of *B. terrestris*. The most robust finding was that instrumental learning and increased required effort were associated with deregulation of genes related to lipid metabolism. Results also suggested a link between depressed expression of mRNA of LOC100646091, encoding cilia and flagella associated protein 91 (CFAP91), to performance of bumblebees in low effort group. However, it is unclear if such relation was due to the task characteristics, or because the low-effort group remained in the same protocol (FR1) for the entire duration of the experiment. Future studies may consider employing a procedure in which subject in the hight-effort group are exposed to increased effort for the whole experimental phase (4 hours). Taken together with our finding it seems reasonable that to learning and effort manipulation in *B. terrestris* during is associated with changes in brain lipid gene expression.

Analyses of the relationship between the top five differentially expressed genes (DEGs) and performance showed that depressed expression of mRNA of LOC100646091 may be associated with low learning performance. In general, CFAP91 protein is a highly evolutionarily-conserved component of motile cilia and flagella. As component of a spoke-associated complex, CFAP91 regulates flagellar dynein activity by mediating regulatory signals between the radial spokes and dynein arms (Bicka et al., 2022). In vitro examination of CFAP91 (LUHMES cells) has shown that down-regulation of CFAP91 may be associated with neurodegeneration (Hollerhage et al., 2022).

Recent observations show that *B. terrestris* queens, workers, and drones differentiate in gene expression (Chen et al., 2024). In addition, previous data suggest that gene expression changes in the bumblebee brain after colour learning may involve changed expression of genes such as Rab10, Shank1 and Arhgap44 (Li et al., 2018). Still, little is known about the link between behaviour and changes in gene expression profile in the cognitive brain and learning in bumblebees foragers. However, evidence exists that pollen diet mediates how pesticide exposure impacts brain gene expression in nest-founding bumblebee queens (Costa et al., 2022). Also, gene expression and related epigenetic processes in pre-imaginal caste differentiation in the primitively eusocial bumblebee *B. terrestris* has been described (Zhu et al., 2021).

Most studies examining the link between neuroplasticity and gene expression profile have, however, been performed using honeybee for example Apis mellifera (Arenas et al., 2021; Iino et al., 2020; Ma et al., 2019). Such studies have earlier suggested that age-based dopaminergic gene expression in the mushroom bodies may be related to behavioural maturation (Humphries et al., 2003). Also, naturally occurring developmental down-regulation of the AChE gene expression has been suggested to facilitate learning capabilities in forager honeybees (Shapira et al., 2001).

In addition new technology that allow for bulk RNA expression, i.e., transcriptomics, has revealed that several EcR target genes may be upregulated in optic lobe glial cells of honeybees during foraging behaviour (Iino et al., 2023). Also, transcriptomic profiles of the mushroom bodies of honeybees may be enriched for GO-terms pointing towards the endoplasmic reticulum in response to nectar or pollen (McNeill et al., 2016). In accordance with these data, the present study showed a trend of enriched GO-terms of endoplasmic reticulum in the high effort reinforcement group. Still, our RNAseq data rather linked the gene expression changes to lipid metabolism.

Previous data show that pheromone exposure has a strong transcriptional effect on the gene expression profile in honeybee foragers (Ma et al., 2019). Moreover, examination of honeybee’s brain for olfactory learning shows that several miRNAs (miR-184-3p, miR-276-3p, miR-87-3p, and miR-124-3p) may be related to olfactory memory (Huang et al., 2023). Recent findings also suggest that down-regulated genes of the brain involved in neurodegenerative processes may result in reduced olfactory discrimination and olfactory sensitivity in failed-learner bees (Raza et al., 2022). In accordance with the observations that down-regulation of some genes may impair learning, the present study suggests that strong down-regulation of LOC100646091 may correlate with low performance as well.

In vitro experiments suggest that honeybee mushroom body D2 type dopamine receptor exhibits age-based plasticity linked to gene expression (Humphries et al., 2003) and that octopamine signalling, which also are related to dopamine brain signalling, affects the decision in honeybee foragers (Arenas et al., 2021). Further evidence shows that expression of Down syndrome cell adhesion molecule (Dscam) – that encodes a molecule required for neuronal wiring – is critically linked to memory retention in adult worker honeybees (Ustaoglu et al., 2024). Taken together, changed expression of such genes may affect plasticity, development, and memory consolidation.

From studies on invertebrates such as C. elegans and insects (e.g., Drosophila) to mammals including mice, lipid metabolism appears to be linked to cognitive functions (Wu et al., 2023). In accordance with these observations, the present study found that neuronal activity during operant learning in B. terrestris is associated with brain energy homeostasis. Moreover, earlier transcriptomic analyses of unconditioned and conditioned behaviour in honey bees have revealed several enriched gene ontology (GO) categories, including “Neuron Differentiation” (Naeger & Robinson, 2016). Thus, gene expression changes in the mushroom bodies associated with operant learning and cognition seem likely. These results are especially relevant and promising, as they provide evidence that instrumental learning through an operant conditioning procedure is linked to genetic expression profiles. This bridges a gap in the literature, as studies evaluating the effects of learning on gene expression typically employ a PER protocol, which investigates reflexive and involuntary learning (Huang et al., 2023).

The two dimensions of behaviour investigated in the present study—instrumental learning and level of effort—may serve as an animal model of flower handling and offer a procedure to understand, for example, the effects of sublethal dosages of toxicants on bee cognition. Only two studies have investigated the effects of pesticides on flower handling, but their procedures were limited in several ways. Stanley and Raine (2016) showed that bumblebees exposed to neonicotinoids performed longer foraging bouts when collecting pollen from novel and morphologically complex flowers; however, their study only measured the duration of foraging bouts and did not identify specific impairments in problem-solving processes. Phelps et al. (2020) investigated the impact of pesticides on flower handling by employing an experimental setup consisting of a small tube that required bumblebees to turn their bodies to reach a sugar-water solution. Although their study provided valuable insights, it had several procedural limitations. The bumblebees’ handling performance was measured as the latency to solve the task, thereby limiting the evaluation of handling over extended periods. Furthermore, their procedure did not allow for manipulation of task difficulty, and the findings were not directly comparable to other studies. The procedure described here involves the learning of a complex instrumental task, with reward delivery manipulated based on the number of lever presses on an operandum. This allows for measurement of bumblebees’ effort and motivation in a task that is both complex (i.e., collecting syrup by activating an operandum) and standardized. Effort-related schedules that manipulate response requirements have been extensively used in the field of behavioral pharmacology to study the effects of drugs on effort and motivation. For example, studies have demonstrated that rats depleted of dopamine exhibit lower breakpoints when required to work for rewards (Salamone et al., 2012).

In the present study, we investigated neuronal changes by focusing on gene expression profiles and demonstrated that exposure to low- and high-effort reinforcement procedures may deregulate numerous genes related to lipid metabolism. Hence, we propose that the procedure used here— learning an operant task and increasing the effort required (number of lever presses), including RNA profiling—may serve as a future model for bumblebee flower handling.

## Author contributions

KC, RS, LE, ES and JG designed the research. RS run the behavioural experimental sessions and L did the brain dissection. KC and RS in discussion with JG analysed the behavioural data. JG performed the genetic data and performed analyses in discussion with KC and RS. KC, RS and JG wrote the manuscript with comments from LE and ES.

## Acknowledgements

We thank Andrine Risøy for the excellent technical support.

---

**Table S1.**
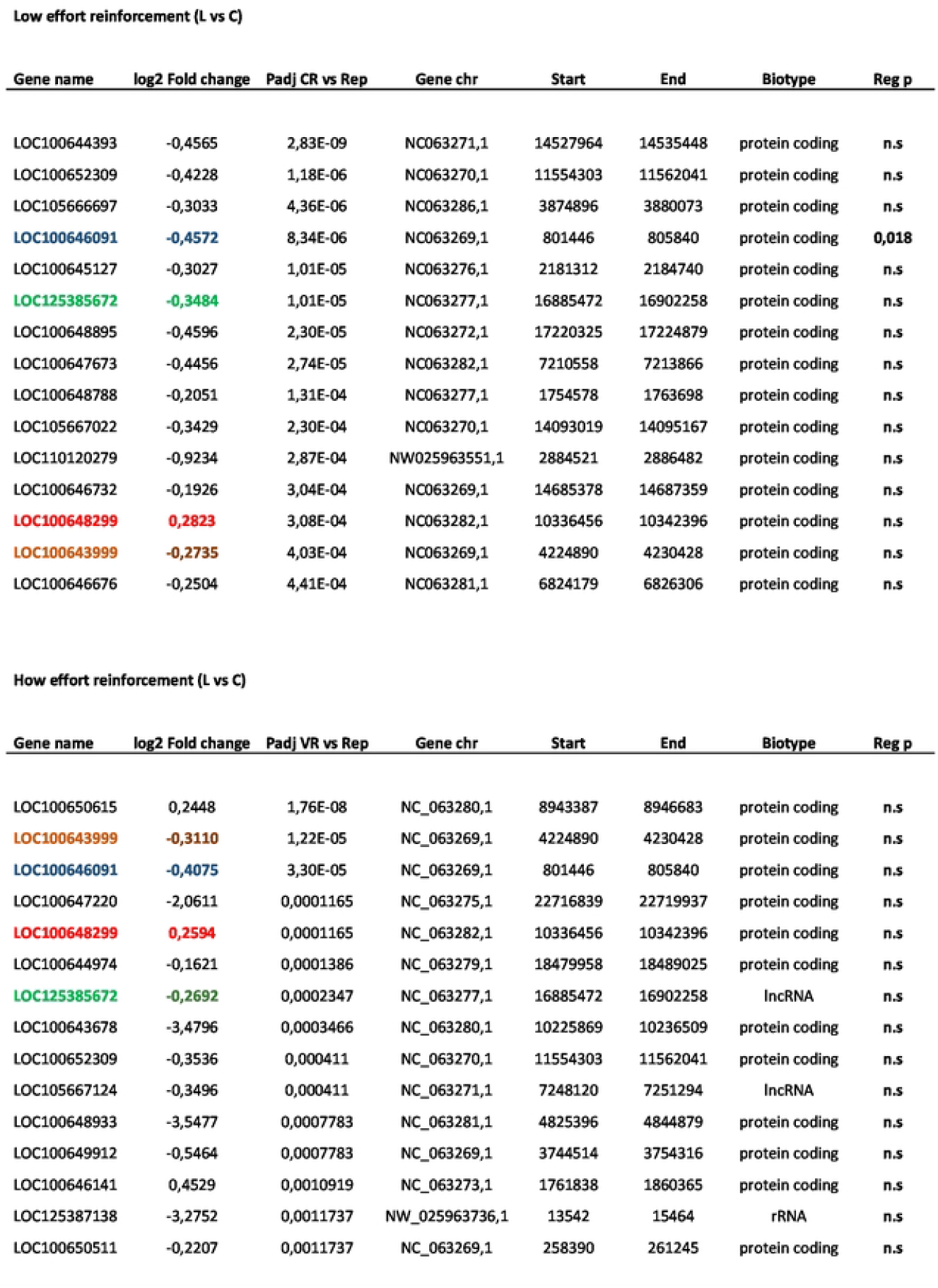
Top 15 differentially expressed genes (DEGs) ranked by adjusted p values (padj) in low effort reinforcement (L) and high effort reinforcement (H) groups relative to control (C) group. Genes found on both lists are highlighted by colours (blue, green, red and brown). Note the gene seen on top five on both lists: LOC100646091.

**Low effort reinforcement (L vs C)**
| Gene name | log2 Fold change | Padj CR vs Rep | Gene chr | Start | End | Biotype | Reg p |
| --- | --- | --- | --- | --- | --- | --- | --- |
| LOC100644393 | -0,4565 | 2,83E-09 | NC063271,1 | 14527964 | 14535448 | protein coding | n.s |
| LOC100652309 | -0,4228 | 1,18E-06 | NC063270,1 | 11554303 | 11562041 | protein coding | n.s |
| LOC105666697 | -0,3033 | 4,36E-06 | NC063286,1 | 3874896 | 3880073 | protein coding | n.s |
| **LOC100646091** | **-0,4572** | 8,34E-06 | NC063269,1 | 801446 | 805840 | protein coding | **0,018** |
| LOC100645127 | -0,3027 | 1,01E-05 | NC063276,1 | 2181312 | 2184740 | protein coding | n.s |
| **LOC125385672** | **-0,3484** | 1,01E-05 | NC063277,1 | 16885472 | 16902258 | protein coding | n.s |
| LOC100648895 | -0,4596 | 2,30E-05 | NC063272,1 | 17220325 | 17224879 | protein coding | n.s |
| LOC100647673 | -0,4456 | 2,74E-05 | NC063282,1 | 7210558 | 7213866 | protein coding | n.s |
| LOC100648788 | -0,2051 | 1,31E-04 | NC063277,1 | 1754578 | 1763698 | protein coding | n.s |
| LOC105667022 | -0,3429 | 2,30E-04 | NC063270,1 | 14093019 | 14095167 | protein coding | n.s |
| LOC110120279 | -0,9234 | 2,87E-04 | NW025963551,1 | 2884521 | 2886482 | protein coding | n.s |
| LOC100646732 | -0,1926 | 3,04E-04 | NC063269,1 | 14685378 | 14687359 | protein coding | n.s |
| **LOC100648299** | **0,2823** | 3,08E-04 | NC063282,1 | 10336456 | 10342396 | protein coding | n.s |
| **LOC100643999** | **-0,2735** | 4,03E-04 | NC063269,1 | 4224890 | 4230428 | protein coding | n.s |
| LOC100646676 | -0,2504 | 4,41E-04 | NC063281,1 | 6824179 | 6826306 | protein coding | n.s |

**How effort reinforcement (L vs C)**
| Gene name | log2 Fold change | Padj VR vs Rep | Gene chr | Start | End | Biotype | Reg p |
| --- | --- | --- | --- | --- | --- | --- | --- |
| LOC100650615 | 0,2448 | 1,76E-08 | NC_063280,1 | 8943387 | 8946683 | protein coding | n.s |
| **LOC100643999** | **-0,3110** | 1,22E-05 | NC_063269,1 | 4224890 | 4230428 | protein coding | n.s |
| **LOC100646091** | **-0,4075** | 3,30E-05 | NC_063269,1 | 801446 | 805840 | protein coding | n.s |
| LOC100647220 | -2,0611 | 0,0001165 | NC_063275,1 | 22716839 | 22719937 | protein coding | n.s |
| **LOC100648299** | **0,2594** | 0,0001165 | NC_063282,1 | 10336456 | 10342396 | protein coding | n.s |
| LOC100644974 | -0,1621 | 0,0001386 | NC_063279,1 | 18479958 | 18489025 | protein coding | n.s |
| **LOC125385672** | **-0,2692** | 0,0002347 | NC_063277,1 | 16885472 | 16902258 | lncRNA | n.s |
| LOC100643678 | -3,4796 | 0,0003466 | NC_063280,1 | 10225869 | 10236509 | protein coding | n.s |
| LOC100652309 | -0,3536 | 0,000411 | NC_063270,1 | 11554303 | 11562041 | protein coding | n.s |
| LOC105667124 | -0,3496 | 0,000411 | NC_063271,1 | 7248120 | 7251294 | lncRNA | n.s |
| LOC100648933 | -3,5477 | 0,0007783 | NC_063281,1 | 4825396 | 4844879 | protein coding | n.s |
| LOC100649912 | -0,5464 | 0,0007783 | NC_063269,1 | 3744514 | 3754316 | protein coding | n.s |
| LOC100646141 | 0,4529 | 0,0010919 | NC_063273,1 | 1761838 | 1860365 | protein coding | n.s |
| LOC125387138 | -3,2752 | 0,0011737 | NW_025963736,1 | 13542 | 15464 | rRNA | n.s |
| LOC100650511 | -0,2207 | 0,0011737 | NC_063269,1 | 258390 | 261245 | protein coding | n.s |

